# High density invasion by the shrub *Ardisia crenata* in Florida, USA is associated with altered soil chemical properties and microbial communities

**DOI:** 10.1101/2025.09.18.676188

**Authors:** Naoto Nakamura, Hirokazu Toju, S. Luke Flory, Kaoru Kitajima

## Abstract

Invasive plants can reshape soil ecosystems by altering chemical conditions and microbial communities. However, field evidence for how these effects scale with invader density, and whether responses are nonlinear, remains limited. We examined whether local stem density of the invasive shrub *Ardisia crenata* is associated with shifts in soil chemistry and soil microbiomes. The local density of *A. crenata*, which reproduces clonally with apomictic seeds, forms a gradient from high density to low density across invaded stands. Our results show strong differences in soil ecosystems with invasion, including consistently lower soil moisture, pH, organic matter, total carbon, total nitrogen, and electrical conductivity. These changes were accompanied by shifts in the community composition of soil prokaryotes and fungi, such as increased abundances of *Acidothermus* and *Saitozyma* with higher density of *A. crenata*. While the exact mechanisms causing these changes could not be established, increases of more labile above- and below-ground litter, as well as *A. crenata* specific secondary metabolites, were possible causal factors. We conclude that strong changes in soil ecosystems can be caused by invasive plants in a mesic forest understory, especially when they occur at high densities. Hence, to maintain healthy soil ecosystems, it is recommendable to control aggressive invasive plants such as *A. crenata* before they reach high densities.

## Introduction

Invasive plants can substantially alter ecosystems, potentially degrading ecosystem services (Kumschick et al. 2015). Competitive dominance by an invasive plant modifies biotic and abiotic environments of the soil (Rodrigues et al. 2015, Anthony et al. 2020), often through input of root exudates and leaf litter that can alter soil chemical properties and microbial communities (Ehrenfeld et al. 2001, Wolfe and Klironomos 2005, Kulmatiski et al. 2008, Stefanowicz et al. 2016, Ahmad Dar et al. 2023). Yet, assessing their impacts in the field remains challenging. Comparisons between invaded and uninvaded sites often confound invasion levels with background variability in soil properties (Stefanowicz et al. 2017, Hussain et al. 2023). Thus, effects of plant invasions on soil chemical properties and microbial community may not be clearly separated, possibly leading to mixed results among studies. Time since invasion, degree (i.e. local density) of invading plants, and site-specific factors, in addition to traits of the invading plant species and the invaded sites, can influence results of studies on how invasions modify soil ecosystems (Sofaer et al., 2018).

As the local density of an invasive plant species increases, so does the supply of litter and root exudates from that species, resulting in cumulative influences on soil microbiomes with time since invasion. Many studies substitute time with space within the same type of background environment (e.g., Gibbons et al., 2017 for a grassland ecosystem). During the active phase of invasion, heterogenous degrees of dominance exist from the original point of colonization (very high local density) to the edge of expanding population (low to negligible presence) (Ren et al. 2022). This is the case for our study species, *Ardisia crenata*as detailed later. However, the relationship between the local density of invasive plant and its impacts on ecosystem may be linear or nonlinear (Yokomizo et al. 2009), and greenhouse experiments manipulating plant density have demonstrated nonlinear impacts on both the soil microbiome and soil chemical properties (Zhang et al. 2020a). Nevertheless, pot experiments may not adequately simulate natural ecosystems, where complex spatial and environmental factors can strongly influence soil microbiomes (Bahram et al. 2018). Hence, it is necessary not only to compare sites with and without invasion, but to consider invader densities in the field, which may be useful for developing effective management strategies of invasive plants, if soil properties are altered beyond recovery once a certain density is exceeded (Schooler et al. 2006, Wu and Carrillo 2016).

In natural ecosystems, soil microbes play significant roles in nutrient cycling, provisioning of essential plant nutrients (Trognitz et al. 2016), and interactions with each other and with plants (Bever et al. 2010). These complex interactions can be disrupted by invasions of specific plant species, which result in either enrichment or depletion of specific bacterial or fungal taxa. If such changes in the soil microbiome establish positive plant–soil feedbacks, they enforce the dominance of invasive species (Anthony et al. 2017, Rodríguez-Caballero et al. 2020, Roche et al. 2023). For instance, *Alliaria petiolata* releases glucosinolates that act as allelopathic compounds, which suppress arbuscular mycorrhizal fungi and reshape the soil microbial community (Anthony et al. 2017). Plant invasion can also alter soil chemical properties, such as pH, nutrient availability and organic matter content through root exudation and litter deposition, and their influence on microbial communities (Ehrenfeld 2003, Inderjit and van der Putten 2010). An increase in invasive plant density should correlate with the degree of enrichment or depletion of specific soil microbial taxa and chemical compounds in the soil. Clarifying how stem density affects soil microbiomes will contribute to a better understanding of soil modification and its subsequent impact on plant-soil feedbacks in the context of plant invasion.

*Ardisia crenata* is a shade-tolerant evergreen shrub that is native to East Asia. It has become widely known for its aggressive invasion of mesic forest understories in the southeastern United States (Dozier 2000, Kitajima et al. 2006, Nakamura et al. 2023). It is an ornamental and medicinal plant that contains triterpenoid saponins in its root (Jia et al. 1994). In Florida, it can create a local density as high as 300 individuals per square meter (Kitajima et al. 2006). Initial colonization in forest ecosystems may occur via animal-mediated seed dispersal or through horticultural introductions in adjacent areas. Recently, it was shown that *A. crenata* individuals consist of a few genetic clones produced by apomixis in Florida (non-native range) and Honshu, Japan (considered to be a native range, but more likely naturalized populations that escaped cultivation) (Noyori et al. 2024). Clonal reproduction via apomictic seeds allows invasion by a single seed and subsequent population growth without another individual. Seed germination level is high (near 100%), and a young plant can start flowering when it is above 20 cm in height. Seeds dropped in the vicinity to create dense patches of *A. crenata*, which continue to expand within a mesic forest understory. Thus, within a given site, the local density of *A. crenata* tends to be the highest at the original point of invasion, with gradual decreases with distance. This pattern results in a clear population density gradient over a relatively small spatial scale of tens to a few hundred meters, making *A. crenata* an ideal model for exploring how the local density of an invasive shrub might modify the soil within the same background habitat.

In this study, we quantified local stem density of *A. crenata*, and distinguished three levels of invasion within a relatively homogeneous forest understory: (1) no *A. crenata* but only native plant species (uninvaded), (2) a mixture of *A. crenata* and native species (low density), and (3) very high dominance of *A. crenata* (high density). We then compared the diversity and structure of soil prokaryotic and fungal communities with high-throughput Illumina sequencing, along with soil chemical properties across the three invasion levels. Through this comparison, we aimed to examine how the variation in the local stem density of *A. crenata* is accompanied by shifts in soil microbial communities and chemical properties.

## Materials and Methods

### Field site

We selected 0.85 ha in 2022 that was experiencing active invasion by *A. crenata* in a mesic hardwood forest in Alachua County, Florida, USA (Figure 1). According to the property owner, *A. crenata* invasion became noticeable about 15 years prior to this study. Whereas active efforts to remove *A. crenata* have been conducted on public lands within 5-km of this site, no such efforts to remove *A. crenata* manually or with herbicide application had been done at our study site. Mean annual temperature is 20.7 D, and mean annual precipitation is 1227 mm for Alachua County (1991-2020, Florida Climate Center). The terrain sloped gently towards a small creek draining to a wetland. The elevation difference was less than 1 m within the site, although it was not precisely determined with a survey instrument. Canopy species within the 0.85 ha stand included *Liquidamber styraciflua, Carya glabra, Magnolia grandiflora, Tilia americana, Quercus nigra*, and *Quercus hemisphaerica*.

**Figure 1.**
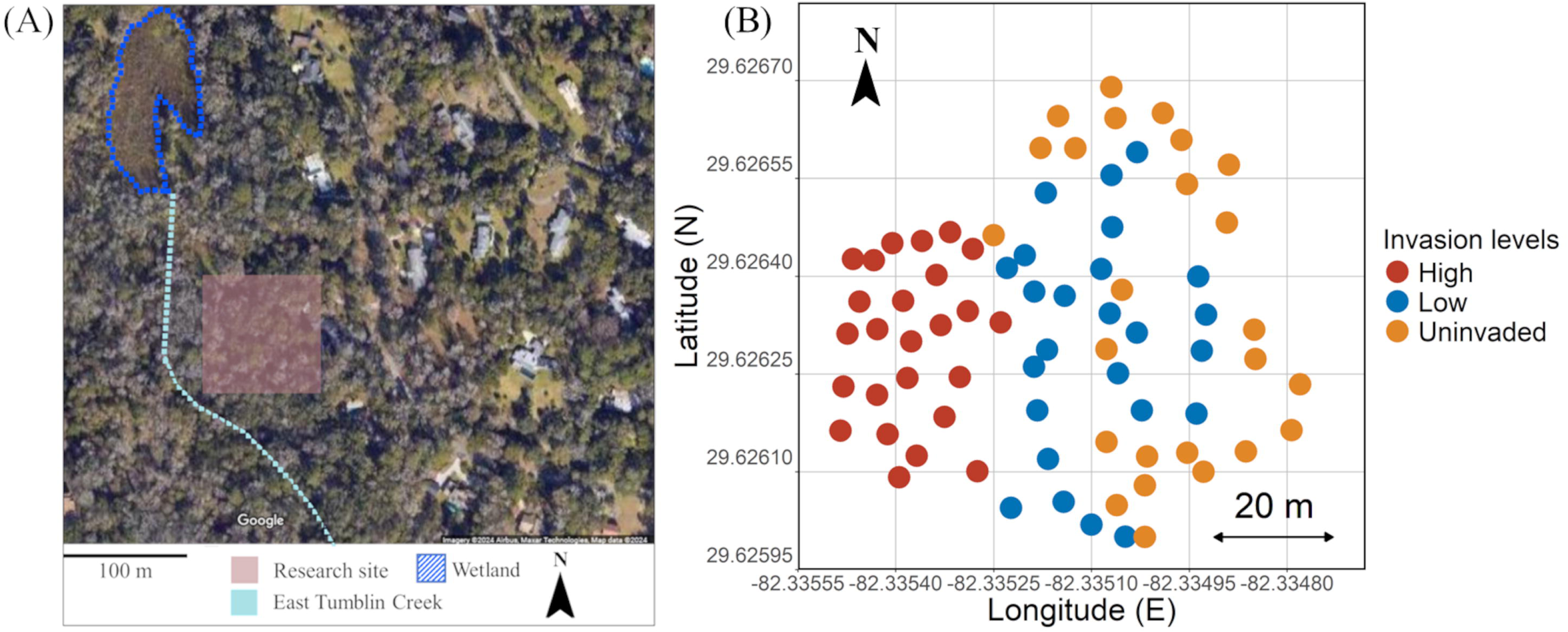
Maps of the research site in Alachua County, Florida. (A) Sky view of the landscape obtained from Goole Map (download date: 2024.11.21) surrounding (B) the study area, in which 75 circular plots (1-m^2^ each, colored dots) were set to encompass three levels of invasion by *A. crenata* (high-density, low-density, uninvaded, indicated by different colors). In (A), the reddish rectangle represents 0.9 ha research site, the location of a wetland is indicated by the blue dotted border, to which a creek (the light blue line) flows into. In the right panel, each circle represents sampling plots.

A total of seventy-five 1 m^2^ circular plots were established. Additionally, to verify whether local stem density measured in 1-m² plots reliably represents surrounding environmental conditions, we also assessed *A. crenata* stem density in the extended 3-m² plots centered on the 1-m² plots. Local density was determined by the number of *A. crenata* individuals taller than 20 cm (adults) within each 1-m² plot (Figure 1). For this study, we distinguished three invasion levels: uninvaded (no adults within 1 m^2^), low density (from one to five adults within 1 m^2^), and high density (more than five *A. crenata* adults within 1 m^2^). For high- and low-density plots, the tallest adult within the plot was chosen as the “focal individual”, and the plot was set to have it at the center. Each plot was placed at a minimum distance of 3 meters from any neighboring plot. Within all high-density plots, the cover of *A. crenata* ranged 70 – 100% (mean = 91.2%). In low-density plots and uninvaded plots, understory vegetation (within 1 m above the ground) included native plants such as *Lindera benzoin, Parthenocissus quinquefolia, Smilax glauca, Acer negundo* and *Toxicodendron radicans*, as well as some non-native plants such as *Tradescantia fluminensis, Dioscorea bulbifera, Tetrapanax papyrifera*, and *Agerantina altissima*.

### Soil sampling and aboveground survey

At each plot, we collected two soil samples using a core sampler (5 cm diameter, 5 cm depth). These two samples were then homogenized to create one composite sample per plot. For high- and low-density plots, samples were taken at two points within 20 cm from the base of the focal individual. For uninvaded plots, soil cores were sampled near the center of the plot. Within a few hours after collection, each composite soil sample was sieved (< 2 mm). A small subsample was sealed in plastic bags with silica gel for drying for about a week, then stored at -20 D until DNA extraction. The remaining portions of the soil samples were air-dried at room temperature for chemical analysis.

We documented the vegetation within each 3-m² plot centered around each 1-m² plot, recording all plant species, the degree of canopy openness at 1.5 m above the ground, the stem number of *A. crenata* adults, and the cover of *A. crenata* and other plant species (estimated as < 1% or in 10% increments from 10% to 100%). The degree of canopy openness was measured as an index of light intensity on the forest floor by taking hemispherical images with RICOH Theta SC2 (RICOH, Tokyo). The detailed methodologies for calculating canopy openness followed ter Steege (2018). Soil moisture was estimated at three randomly selected points within each 1 m^2^ plot using HH1 Theta Probe (Organic soil moisture mode; Delta-T Devices, Cambridge, England).

### Soil chemical properties

Soil organic matter content was measured by the loss-on-ignition (LOI) method, with samples combusted at 550 °C for 12 hours. Total nitrogen and total carbon concentrations were analyzed using a Flash 2000 CN soil analyzer (Thermo Fisher Scientific, USA). For soil pH and electrical conductivity (EC) measurements, a 1:5 (weight/volume) suspension of air-dried soil and deionized water was prepared. Both soil pH and EC were then measured from this suspension using an Electrode GST-5721C pH meter (TOA-DKK, Japan).

### DNA extraction, PCR amplification, and sequencing

Soil microbial DNA was extracted from 0.25 g of soil using the DNeasy PowerSoil DNA Isolation kit (Qiagen, Hilden, Germany) following the manufacturer’s protocol with some modifications. The extracted DNA was dissolved in TE buffer and stored at −20°C until further processing. A negative control was included during DNA extraction: 0.25 mL of sterile water was processed instead of a soil sample using the same method to confirm the absence of contamination.

We used a two-step PCR approach for Illumina MiSeq library preparation. The prokaryotic 16S rRNA and fungal internal transcribed spacer 1 (ITS1) regions were PCR amplified following the protocol detailed elsewhere (Toju et al. 2019) with some modifications. Briefly, for the prokaryotic 16S rRNA region, the primer set 515f/806rB (515f, Caporaso et al., 2011; 806rB, Apprill et al., 2015) was used. Likewise, the fungal ITS1 region was amplified using the primer set ITS1-F_KYO1/ITS2_KYO2 (Toju et al. 2012). Both primer sets were fused with the Illumina sequencing adapter sequences and 3-6-mer Ns to improve sequencing quality (Lundberg et al. 2013). For both 16S rRNA and ITS1 regions, first PCR was performed using Platinum Super D (Invitrogen, USA) under the following thermal cycling conditions of initial denaturation 98°C 30 seconds, 35 cycles at 98 °C for 10 seconds (denaturation), 65 °C for 30 seconds (annealing of primers), and 72 °C for 30 sec (extension), and a final extension at 72 °C for 5 min. Amplification of the DNA fragments was confirmed using the Flash Gel System for DNA (Lonza, Switzerland). The PCR products were purified using the AMPureXP Kit (Beckman Coulter, USA) to remove primer dimers. The ratio of AMPureXP reagent to each of the first PCR products was set to 0.5 (v/v). We added a negative control (PCR negative control) for both 16S rRNA and ITS1 PCR reactions to confirm the absence of contamination.

A second-round PCR was carried out to append indices for different samples for Illumina MiSeq sequencing. PCR was performed using KAPA HiFi HS (KAPA Biosystems, USA) under the following thermal cycling conditions: 35 cycles of 98 °C for 20 s (denaturation), 65 °C for 15 s (primer annealing), and 72 °C for 30 s (extension), followed by a final extension at 72 °C for 6 min for both 16S rRNA and ITS1 samples. The second PCR products were purified using the AMPureXP Kit and quantified using Qubit 4.0 fluorometer (Invitrogen, USA). The ratio of AMPureXP reagent to each second PCR product was set to 0.5 (v/v). For each 16S rRNA and ITS1 library, equimolar amounts of second PCR products were pooled based on quantification results from the Qubit dsDNA HS Assay (Thermo Fisher Scientific, USA). To further remove primer dimers, we performed electrophoresis using E-Gel™ SizeSelect™ II Agarose Gels, 2% (Invitrogen, USA) and excised bands of desired DNA size from the gel. The pooled library was sequenced using an Illumina MiSeq sequencer with 10% PhiX spike-in (Center for Ecological Research, Kyoto, Japan, 2 × 300 cycles).

### Bioinformatics

The raw sequencing data were converted into FASTQ files using the program bcl2fastq 1.8.4 distributed by Illumina. The output FASTQ files were then demultiplexed with Claident v0.2.2018.05.29 (Tanabe and Toju 2013). Illumina adapters were removed using Cutadapt v4.4 (Martin 2011). The sequencing reads were subsequently processed using the package “DADA2” v.1.18.0 of R version 4.3.1 (R Core Team 2022) to remove low-quality sequences with quality score below 20, and to remove chimeras (function ‘FilterTrimming’ and ‘removeBimeraDenovò, respectively). All reads included in the negative controls were confirmed to have been removed at this stage. Molecular identification of the obtained amplicon sequence variants (ASVs) was performed using the naive Bayesian classifier method (Wang et al. 2007) with the ‘assignTaxonomy’ function in DADA2. Specifically, 16S rRNA ASVs were classified using the SILVA v.132 database, and ITS1 ASVs were classified using the UNITE database v9.0 release (Abarenkov et al. 2023). Obtained ASVs were clustered into OTUs at 97% similarity using Vsearch (Rognes et al. 2016) for both prokaryotes and fungi. To generate the final OTUs table, we removed nonprokaryotic and nonfungal OTUs. In total, 5096 prokaryotic OTUs (2,084,997 reads) and 3464 fungal OTUs (1,311,726 reads) were detected.

The dataset was rarefied at 13,521 reads, and 6,314 reads for prokaryotes and fungi, respectively, using the ‘rrarefy’ function of the “vegan” package (Oksanen et al. 2022) in R. Samples that yielded fewer reads than these thresholds were discarded, resulting in 69 prokaryotic and 66 fungal samples. The rarefaction curves for each group reached an asymptote, indicating that our sampling depth was adequate (Supplementary Figure S1). After rarefaction, a total of 4,797 prokaryotic OTUs and 2,334 fungal OTUs were included in the final analysis, respectively.

### Statistical analysis

The relationships between soil properties and the density of *A. crenata* were examined both for 1-m² and 3-m² plots, which showed very similar patterns (Supplementary Figure S2). Some of 1-m^2^ plots that had no *A. crenata* stems were within 3-m² plots with few *A. crenata* stems, but overall, the stem number was highly correlated between 1-m² and 3-m² plots (Pearson’s r = 0.948, p < 0.001). To avoid redundancy, we present the results of local density effects using the data from 1-m² plots. The patterns shown in Figure S2 also justifies the use of three levels of local density of *A. crenata* in the subsequent analyses.

The differences in soil chemical properties and aboveground properties among the three levels of local density of *A. crenata* were tested using the Kruskal-Wallis test followed by Mann-Whitney U post hoc tests for pairwise comparisons (Bonferroni corrected where appropriate). The relationship between observed variables (soil organic matter, pH, electrical conductivity, total carbon, total nitrogen and plant cover) and the number of individuals of *A. crenata* within both 1-m² and 3-m² plots were visualized using scatter plots with quadratic regression. To calculate alpha diversity of prokaryotic and fungal communities, we used the exponent of Shannon diversity (i.e., exp (−ΣPi ln (Pi)); where *Pi* is the proportional abundance of species *i*, Shannon, 1948) using the R package “vegan”. To compare alpha diversity among invasion statuses, we conducted one-way analysis of variance (ANOVA) and a post hoc comparison with a Tukey HSD test (Bonferroni corrected where appropriate) after confirming the normality and homogeneity of variances using a Shapiro-Wilk test and a Bartlett test, respectively. To examine how microbial compositions differed by invasion levels, we conducted permutational multivariate analysis of variance (PerMANOVA, 9,999 permutations; Anderson, 2006) based on Bray-Curtis distance values with the ‘adonis2’ function in “vegan” package. In addition, permutational analysis for multivariate homogeneity of dispersions (PERMDISP; Anderson, 2006) was conducted for bacterial and fungal communities (9,999 permutations). Non-metric multidimensional scaling (NMDS) based on Bray-Curtis dissimilarity was also conducted using “vegan” and “ggplot2” packages (Wickham 2016).

The relative abundance of OTUs at genus, order and phylum levels were visualized as bar graphs, using rarefied data and the “vegan” package in R. Ordination methods were used to determine the relationships between prokaryotic and fungal communities and environmental variables. Detrended correspondence analysis was used as a preliminary step to assess the length of the ordination axis, which was >3 SD (3.46 for prokaryotes and 5.01 for fungi). This pattern suggested that the response curve was unimodal, and that canonical correspondence analysis (CCA) was appropriate (Jongman et al. 1995). The explanatory variables that significantly contribute to the microbial structure were estimated using forward selection with the “vegan” package (Monte Carlo permutation test with 9,999 randomizations). The Linear Discriminant Analysis Effect Size (LEfSe) algorithm was used to identify bacterial and fungal taxa (genus level) that were characteristic of each invasion level (Segata et al. 2011). The Kruskal-Wallis rank sum test was used to detect significantly different abundances and generate LDA scores to estimate the effect size (threshold, 2.5) in the LEfSe analysis. Only genera with greater than 0.1% average relative abundance was used for Lefse analysis, while genera with lower abundance were grouped as “others”.

## Results

### Soil chemical properties

Soil organic matter content, total nitrogen, total carbon, soil pH and electrical conductivity (EC) exhibited significant variations in relation to the local density of *A. crenata* (Figure 2). The overstory cover did not differ significantly among the local density levels of *A. crenata*, but the total understory vegetation cover, which included *A. crenata*, was higher in high-density plots (Table 1). In post-hoc pairwise comparisons, differences between uninvaded plots and low-density plots were significant only for soil C/N ratio, whereas differences between high-density plots and low-density or uninvaded plots were significant for all soil chemical properties except for C/N ratio (Table 1).

**Figure 2.**
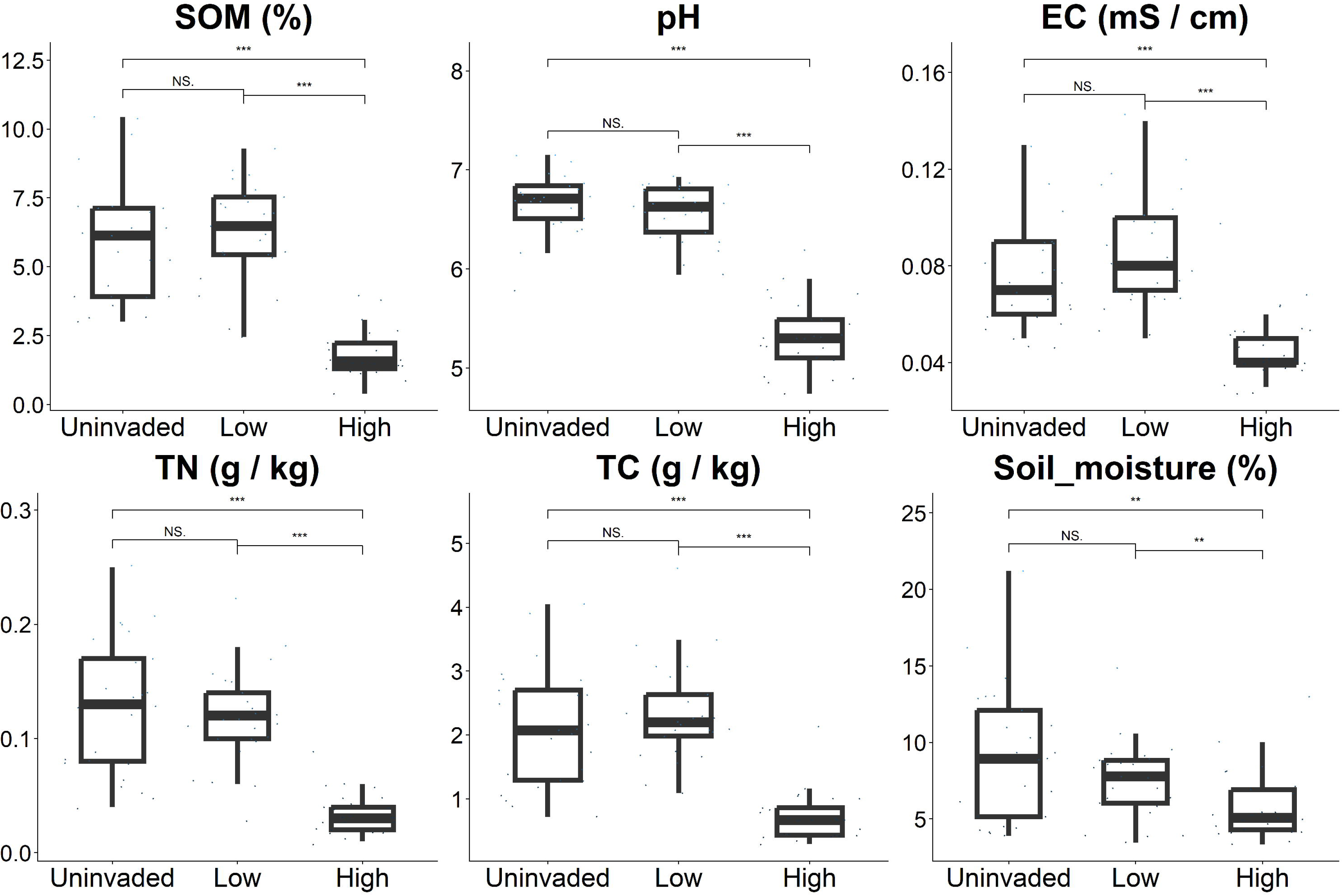
Soil chemical properties summarized by three invasion levels. A significant difference between pairs of invasion levels is indicated by an asterisk, determined by the Mann-Whitney U post hoc tests (\**p* < 0.05, \*\**p* < 0.01, \*\*\**p* < 0.001). SOM: soil organic matter; EC: electric conductivity; TN: total nitrogen; TC: total carbon; Soil moisture; volumetric soil moisture content.

**Table 1.** The result of soil chemical analysis, aboveground survey in three invasion statuses. Mean ± standard deviation (n= 25 each). The degree of freedom and p-value of the result of the Kruskal-Wallis test were shown. Superscript letters indicate the results of pairwise comparison (*p* < 0.05, Mann-Whitney U test with Bonferroni correction).

|  |  | Invasion levels |  |  | Kruskal-Wallis test |  |
| --- | --- | --- | --- | --- | --- | --- |
|  |  | Uninvaded | Low-density | High-density | df | *p* |
| **Soil properties** | Electric conductivity [mS/cm] | 0.07 ± 0.00<sup>a</sup> | 0.09 ± 0.00<sup>a</sup> | 0.05 ± 0.00<sup>b</sup> | 2 | < 0.001 |
| | pH | 6.67 ± 0.06<sup>a</sup> | 6.56 ± 0.06<sup>a</sup> | 5.32 ± 0.07<sup>b</sup> | 2 | < 0.001 |
| | Soil_moisture [%] | 9.08 ± 0.89<sup>a</sup> | 7.54 ± 0.50<sup>a</sup> | 5.85 ± 0.45<sup>b</sup> | 2 | 0.007 |
| | Soil organic matter [%] | 5.96 ± 0.45<sup>a</sup> | 6.25 ± 0.36<sup>a</sup> | 1.82 ± 0.17<sup>b</sup> | 2 | < 0.001 |
| | Total carbon [g / kg] | 2.10 ± 0.18<sup>a</sup> | 2.34 ± 0.15<sup>a</sup> | 0.72 ± 0.08<sup>b</sup> | 2 | < 0.001 |
| | Total nitrogen [g / kg] | 0.13 ± 0.01<sup>a</sup> | 0.12 ± 0.01<sup>a</sup> | 0.04 ± 0.00<sup>b</sup> | 2 | < 0.001 |
| | C / N | 16.9 ± 0.5<sup>a</sup> | 20.8 ± 1.6<sup>b</sup> | 21.0 ± 1.1<sup>b</sup> | 2 | < 0.001 |
| **Aboveground**<br>**Survey** | Canopy_openness [%] | 35.2 ± 2.1 | 39.8 ± 1.8 | 36.5 ± 1.4 | 2 | 0.306 |
| | Plant_cover [%] | 34.8 ± 4.8<sup>a</sup> | 51.2 ± 3.9<sup>b</sup> | 91.2 ± 2.2<sup>c</sup> | 2 | < 0.001 |

### Microbial diversity and community structure

Shannon diversity of prokaryotic communities differed significantly among the three invasion levels of *A. crenata* (ANOVA: *p* < 0.001), with the highest diversity observed in the low-density plots and the lowest in the high-density plots (Figure 3). In contrast, the Shannon diversity of the fungal community did not differ significantly among the three invasion levels of *A. crenata* (ANOVA: *p* = 0.07), although the fungal diversity tended to decrease with increasing stem density of *A. crenata* (Figure 3).

**Figure 3.**
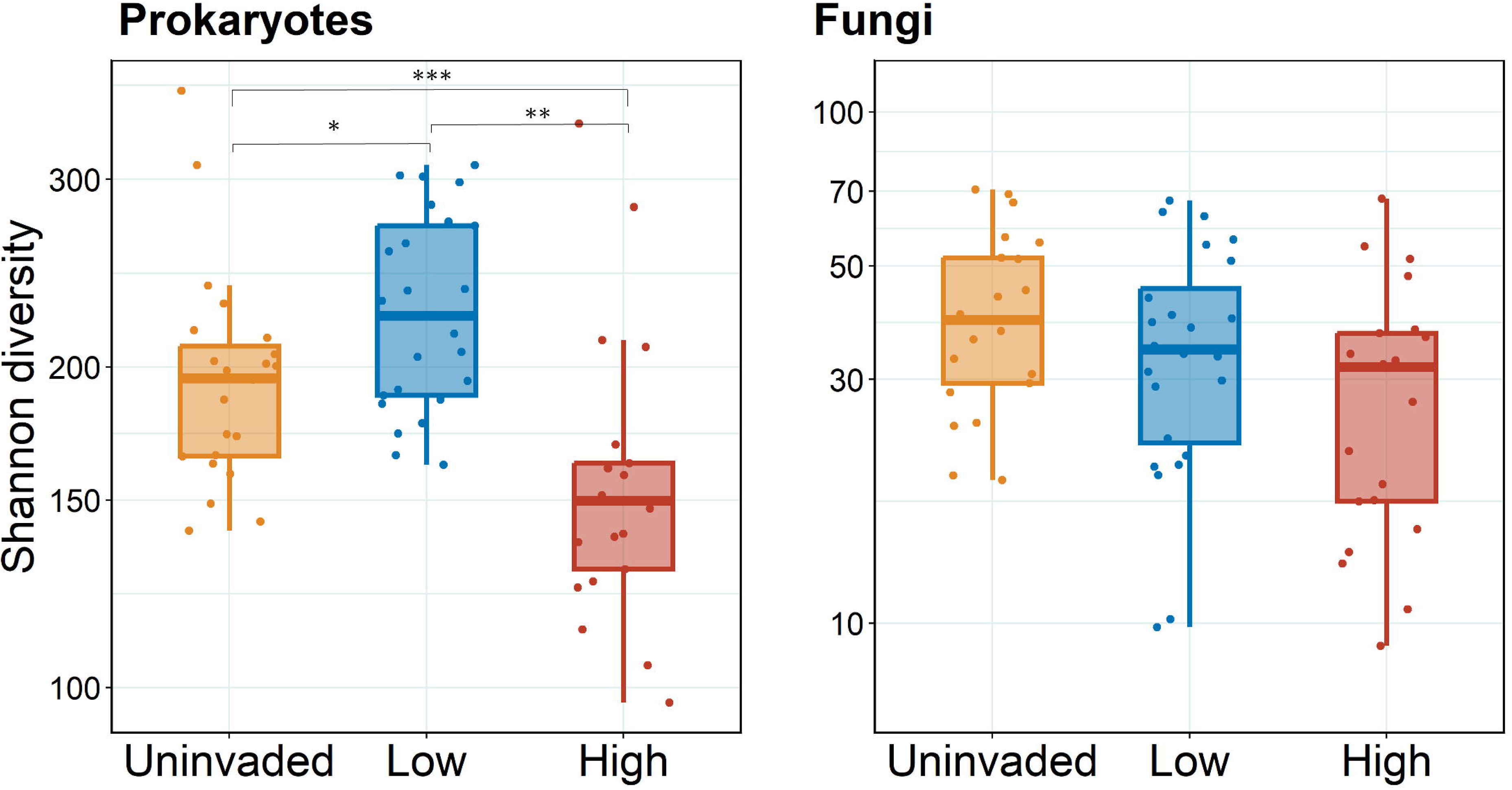
Shannon diversity of prokaryotic and fungal communities at three invasion levels. Significant difference among invasion statuses as indicated by asterisks, determined by the Turkey HSD test (\**p* < 0.05, \*\**p* < 0.01, \*\*\**p* < 0.001).

PerMANOVA showed that invasion levels were significantly related to the prokaryotic and fungal community structures (*p* < 0.001; Supplementary Table S1). For the prokaryotic community, post hoc PerMANOVA showed that the community structure of high-density plots differed significantly from low-density or uninvaded plots, while there were no significant differences between low-density and uninvaded plots (*p* = 0.023). For the fungal community, post hoc PerMANOVA showed that the community structure of high-density plots differed significantly from those of low-density or uninvaded plots, while there was a marginal difference between the latter two types (*p* = 0.001). PERMDISP, which tested whether the heterogeneity of OTU compositions (i.e., dispersion) differed among invasion levels, indicated no significant differences (Supplementary Table S1).

### Taxonomic comparisons in relation to invasion levels

Differences in microbial community composition among the invasion levels were also examined at various levels of taxonomic classification. At the prokaryotic phylum level, soil samples predominantly contained Actinobacteria, Proteobacteria, and Acidobacteriota, which together accounted for approximately 60% of the total abundance irrespective of the invasion levels (Figure 4). In the high-density plots, Actinobacteria and Proteobacteria were more common, while Acidobacteriota was more abundant in the low-density and uninvaded plots. At the order level, Rhizobiales, Vicinamibacterales, Gaiellales, Chthoniobacerales, and Nitrososphaerales were dominant in terms of relative abundance, accounting for approximately 60% of the total abundance in the low-density and uninvaded plots (Supplementary Figure S3). In contrast, the high-density plots had low abundance of Nitrososphaerales and Vicinamibacterales, and greater abundance of Elsterales compared to low-density and uninvaded plots. At the genus level, in particular for bacteria, there were clear differences in genus composition by invasion level,Figure 5,. LefSe analysis identified 21 prokaryotic genera as characteristic of the high-density plots, with *Acidothermus, Conexibacter, and Candidatus Udaeobacter* being prominent genus (Figure 6). In contrast, 11 genera were indicative of the low-density plots, with the *Pir4* lineage as a prominent genus. For the uninvaded plots, 18 genera were characteristic of the low-density areas, with *Gaiella* and *Candidatus Xiphinematobacter* being the most prominent genera.

**Figure 4.**
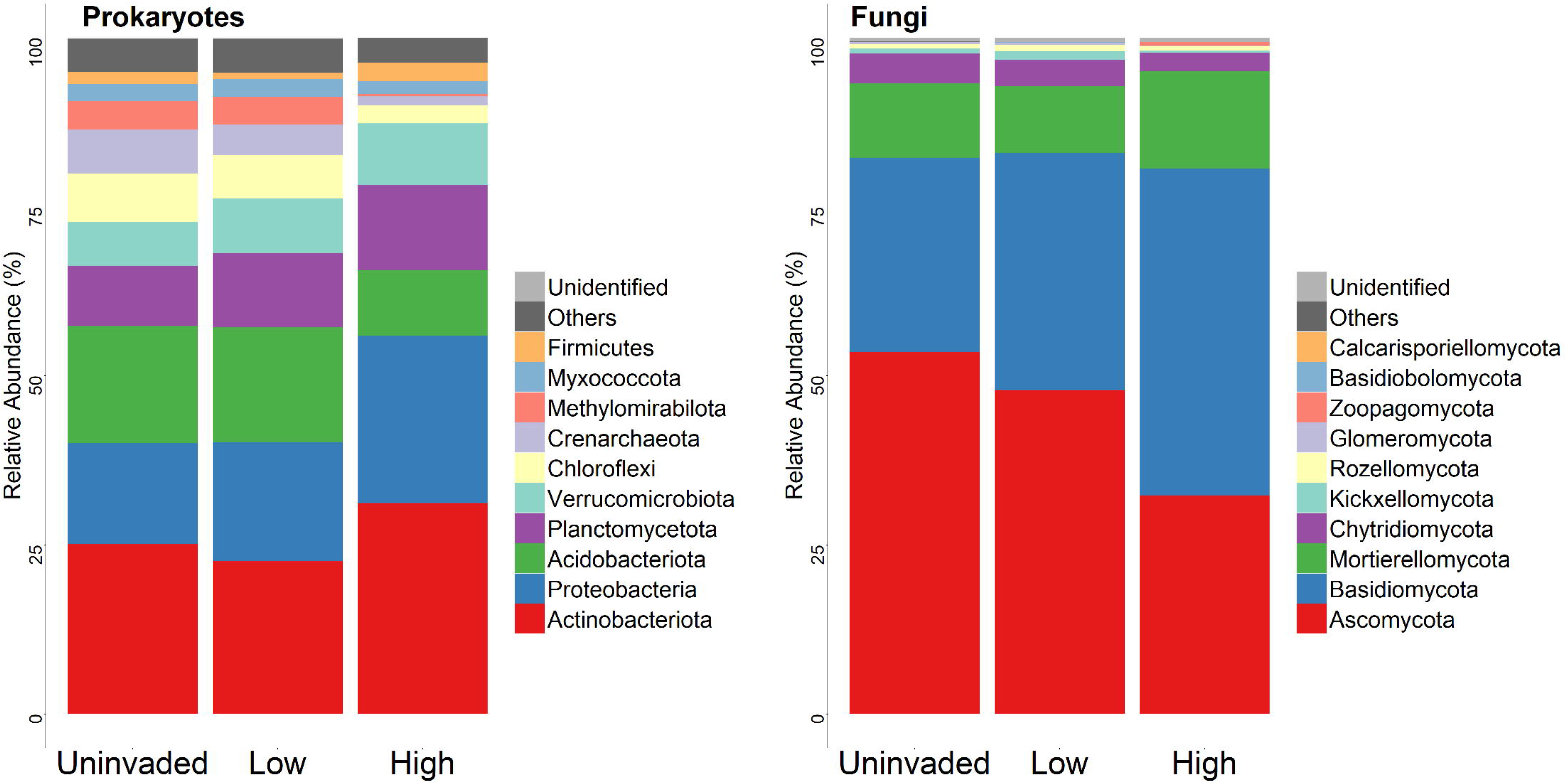
The phylum-level taxonomic composition of prokaryotic and fungal OTUs. The top 10 taxa were displayed. All remaining phyla are consolidated into the ’Others’ category. The vertical axis indicates the relative abundance of each taxon.

**Figure 5.**
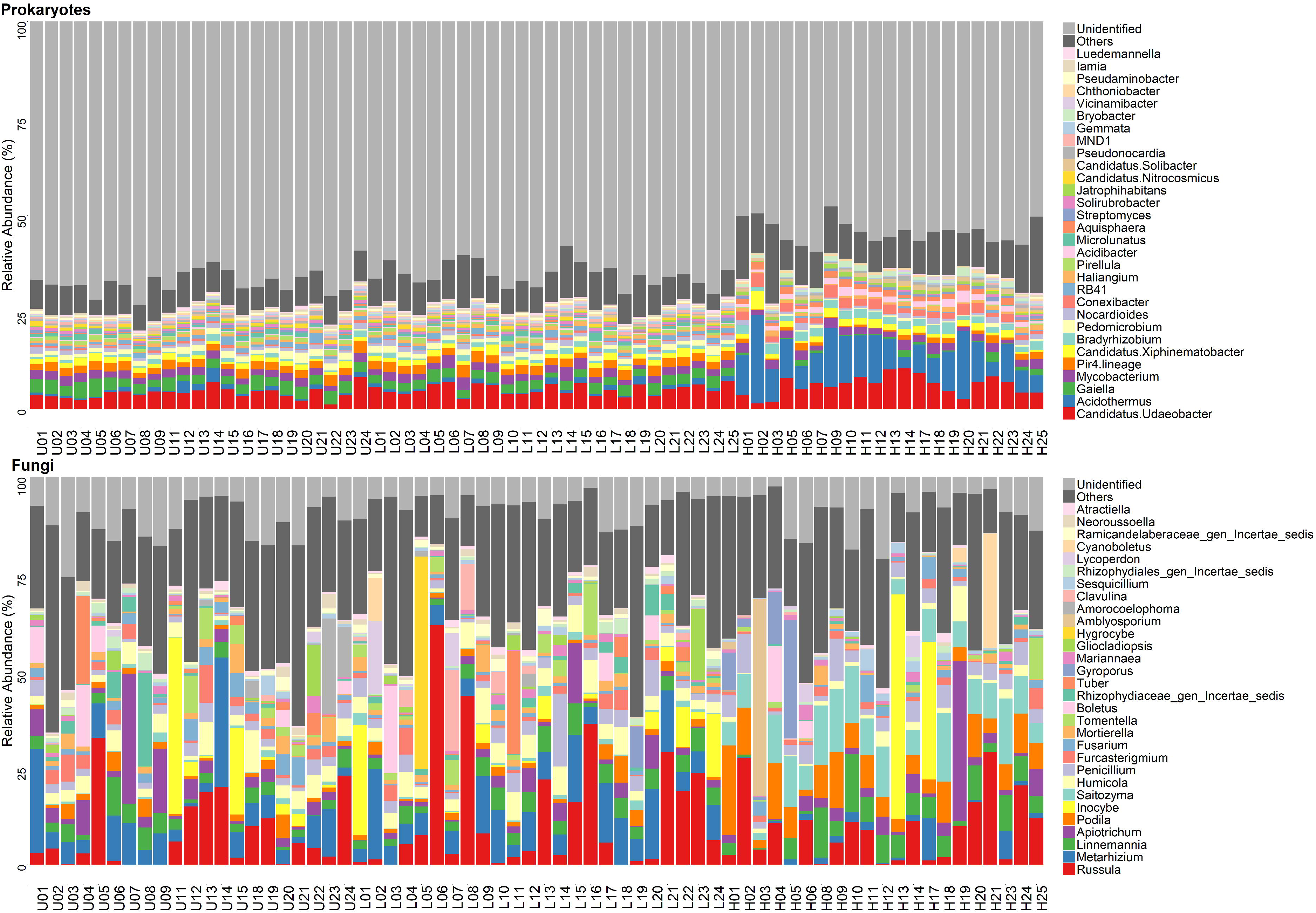
The genus-level taxonomic composition of procaryotic and fungal OTUs in the soil of 75 of 1-m^2^ circular plots. The first letter of the sample ID indicates the invasion levels, H for high-density, L for low-density, and U for uninvaded. The top 30 taxa were displayed. All remaining species are consolidated into the “Others” category. The vertical axis indicates the relative abundance of each taxon.

**Figure 6.**
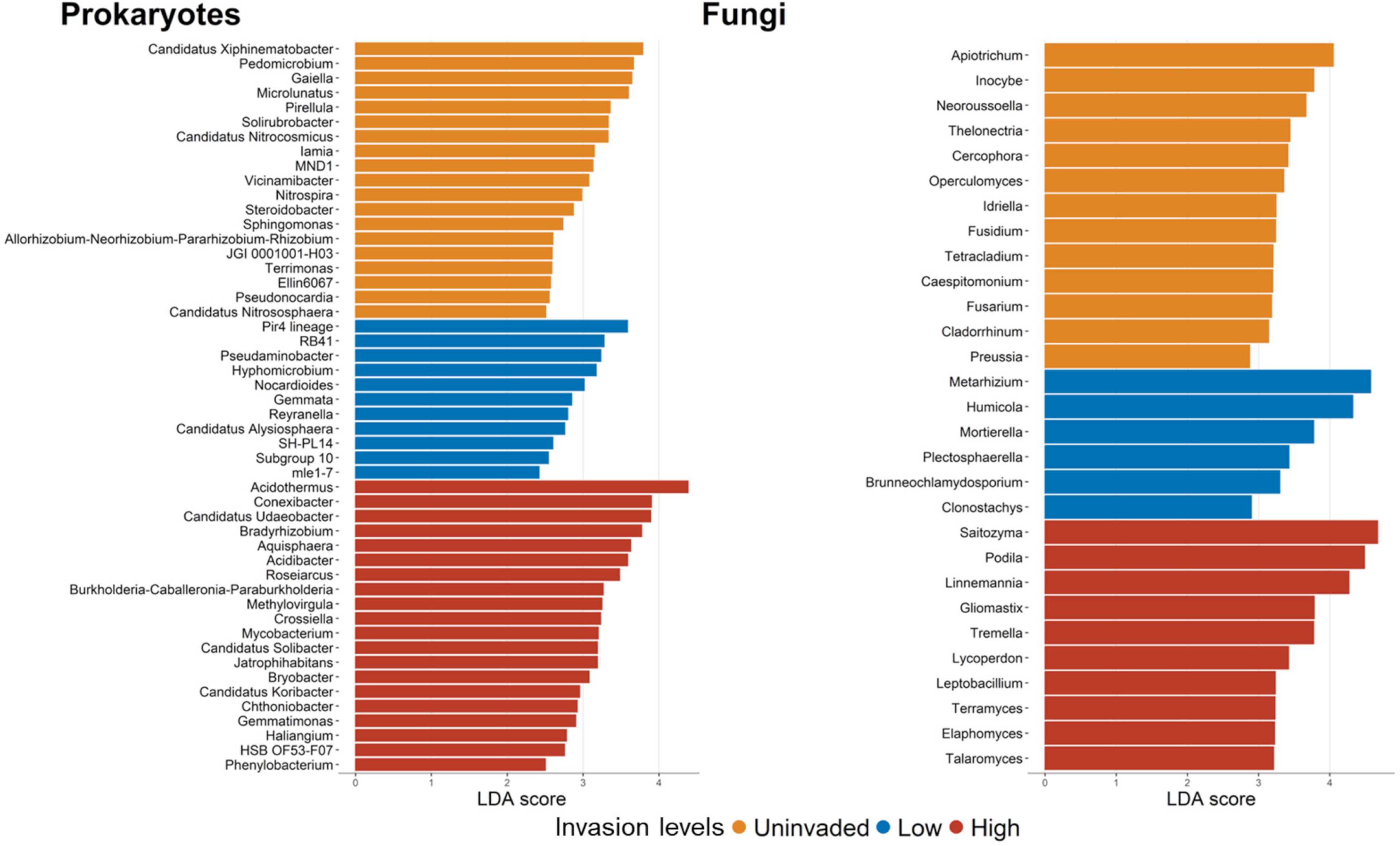
Effect Size of Liner Discriminant Analysis (LDA) for genera of prokaryotes and fungi with LDA > 2.5 at three levels of invasion: uninvaded (yellow), low-density (blue), and high-density (reddish brown) plots.

The fungal community in the soil samples predominantly consisted of Ascomycota and Basidiomycota, comprising about 80% of the total abundance (Figure 4). With increasing local density of *A. crenata*, there was a decrease of Ascomycota and an increase of Basidiomycota. At the order level, Hypocreales and Mortierellales were found in all plots, accounting for approximately 15-30% of the total abundance, with slight tendency of lower abundance of Hypocreales in high-density plots (Supplementary Figure S3). Russulales and Agaricales appeared sporadically in several plots, while Tremellales were more frequently found in the high-density plots, and less so in low-density and uninvaded plots. Conversely, Sordariales were more prevalent in the low-density and uninvaded plots. LefSe analysis identified 11 fungal genera as characteristic of the high-density plots, with *Saitozyma*, *Podila* and *Linnemannia* being the most prominent genera (Figure 6). In contrast, 6 genera were indicative of the low-density plots, with *Metarhizium* and *Humicola* being prominent. For the uninvaded plots, 13 genera were indicative, with *Apiotrichum* and *Inocybe* being the prominent genera.

### Canonical correspondence analysis

For Canonical Correspondence Analysis (CCA) of prokaryotic community, total carbon was removed from the analysis to avoid multicollinearity. The final CCA model, after the forward selection process, included the following variables: pH, organic matter, and moisture. These variables explained 12.7% of the total inertia, with the first two axes accounting for 47.7% and 13.3% of the constrained inertia, respectively (Figure 7). The high-density plots clustered distinctly from low-density and uninvaded plots, and this difference was primarily associated with the first CCA axis, representing a gradient of lower pH and organic matter.

**Figure 7.**
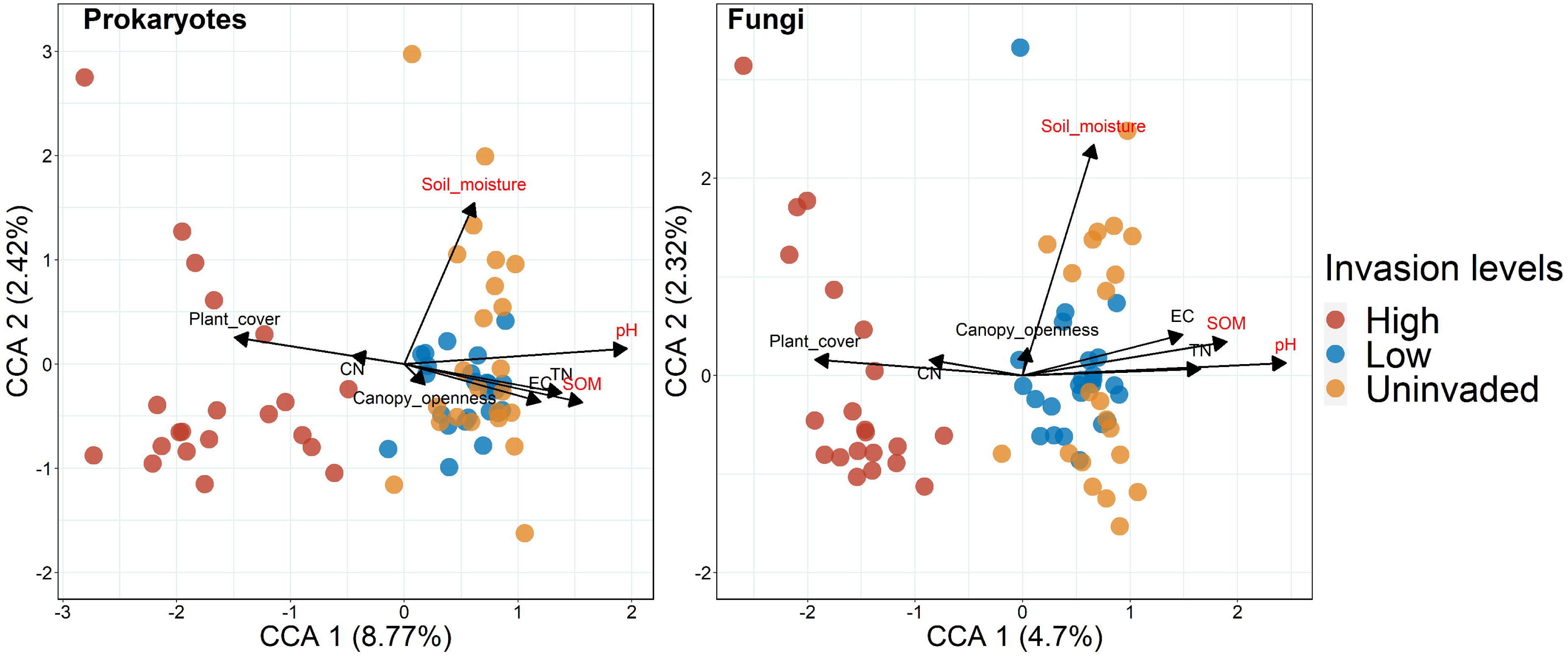
Canonical correspondence analysis (CCA) of soil procaryotes and fungi communities sampled in 1-m^2^ plots at three different levels of invasion: high-density (reddish brown), low-density (blue), and uninvaded (yellow) plots. The dissimilarity matrix at the OTU level is based on Bray-Curtis dissimilarity on relative abundances. Environmental variables are represented as arrows, with key variables influencing community variance highlighted in red labels following a forward selection procedure. Explained proportions of each axis against total inertia are shown. The SOM: soil organic matter; EC: electric conductivity; TN: total nitrogen; Plant_cover: % cover of vegetation including *A. crenata* and others.

For CCA of fungal community, total carbon was similarly removed from the analysis to avoid multicollinearity. The final CCA model, following the forward selection process, included variables: pH, organic matter, and moisture, explaining 8.51% of the total inertia. The first two axes accounted for 46.8% and 23.4% of the constrained inertia, respectively (Figure 7). As in the prokaryotic CCA, high-density plots clustered distinctly in the fungal CCA plots, with its difference primarily explained by the first CCA axis, represented mainly by pH and soil organic matter. Additionally, variations in soil fungal community compositions within each invasion level aligned along the second CCA axis, which was mainly defined by gradients of soil moisture.

## Discussion

We found that soil chemical properties and microbial community structure were modified by high local densities of the invasive shrub *Ardisia crenata*. At low densities, there were limited differences in soil chemical properties or microbial community structure compared to uninvaded plots. This finding suggests that in early invasion stages, the local density of *A. crenata* or the time since site-invasion might be insufficient to induce substantial soil alterations. Similar patterns have been reported from other invasive species. For instance, while the invasion of *Alliaria petiolata* is generally known to affect soil microbiomes (Anthony et al. 2017, 2020), early invasion phases have been reported to show limited effects on microbial communities (Edwards et al. 2022). Our results suggest that once the local density of *A. crenata* becomes high enough (e.g., above 10 stems per m^2^, Figure S2), substantial changes in both soil chemical properties and microbial communities occur. This observed pattern could be attributed to the direct impact of *A. crenata* through litter input and root exudates, although our study did not examine these potential mechanisms.

Our study used space-for-time substitution, based on observations by local researchers that the density of *A. crenata* increases over time from the point of original invasion. Because our results are fundamentally associations, it is possible that the observed differences in soil observed in high density plots from uninvaded or low-density plots might reflect pre-existing soil characteristics that favored high *A. crenata* density in those locations. However, *A. crenata* is known to prefer moist (but not inundated) soils near the edges of rivers and lakes even though it can also colonize drier slopes and ridges (Kitajima et al. 2006). In CCA, soil moisture was not strongly associated with the invasion levels. Although the high-density plots were positioned in low-lying areas adjacent to a creek, they nevertheless had lower soil moisture than plots at higher elevations. Hence, this pattern is unlikely be established by the habitat preference of *A. crenata* for moist habitats. Instead, the observed lower soil moisture in the high-density plots might be caused by high water use by *A. crenata*. However, it would require additional evidence to conclude whether the variation in soil characteristics observed across invasion levels was induced by *A. crenata* or instead reflected pre-existing differences in environmental conditions. Other studies caution that it is difficult to decipher cause vs. effect in observed associations between plant invasion and soil properties (Gibbons et al. 2017, Rodríguez-Caballero et al. 2020).

There were no apparent differences in overstory tree composition within our study site. If our results indeed indicate that an understory shrub modifies soil chemical properties and microbial communities, this would be consistent with previous findings that understory vegetation can also play a significant role in shaping soil properties (Zhao et al. 2014). The observed non-linear relationship between the local density of *A. crenata* and soil characteristics (Figure S2) is consistent with previous reports on invasive-plant impacts, including density-dependent effect of invasive grasses (Gibbons et al. 2017, Frost et al. 2024) and non-linear effects of invasive *Conyza canadensis* on soil characteristics in a pot experiment (Zhang et al. 2020a). Our results, along with these other studies, strongly suggest the presence of density thresholds at which invasive plants can significantly alter both biotic and abiotic soil properties (Foxcroft 2009). Once an invasive species reaches a high density and alters the biotic and abiotic factors of the soil, its removal may not easily reverse these changes (Sofaer et al. 2018). These findings underscore the importance of early intervention before the local density of *A. crenata* becomes very high, as post-threshold changes may be difficult or impossible to reverse.

Among the soil chemical properties, marked changes were observed in soil organic matter content (SOM) and total nitrogen (TN), both of which were lower in high-density *A. crenata* plots. This finding contrasts with the general trend reported in meta-analyses, which show that plant invasion often increases SOM and TN due to enhanced plant biomass and soil enzymatic activity (Xu et al. 2022). Because *A. crenata* is an understory shrub, the leaf litter input may be small relative to the leaf litter input from overstory trees. Thus, it is more likely that the observed decrease in SOM reflected enhanced litter decomposition rates. In a mesic forest in Florida, the same type of forest as used in our study, Bray et al. (2012) compared litter chemistry and decomposition of common plant species, including *A. crenata*, which was found to be intermediately labile to decompose (4^th^ among the 10 species). Studies with other species have reported that invasive plants accelerate SOM decomposition rates via microbial priming and/or supply of labile litter (Allison and Vitousek 2004, Tamura and Tharayil 2014). *A. crenata* also produces roots with high concentrations of non-structural carbohydrates (Kitajima et al. 2006). Although further experimental investigation is needed, it is possible that microbial communities had changed in response to greater input of labile above- and below-ground litter in the high-density plots dominated by *A. crenata*, which accelerated overall decomposition rates.

The small but significant increase in Shannon diversity of soil prokaryotes from uninvaded to low-density plots may reflect a higher overall understory vegetation cover and/or greater labile litter input. Others report that the addition of non-native plants can enhance plant community diversity, which in turn may positively affect the diversity of plant-associated microbes (Lu-Irving et al. 2019). The lower Shannon diversity observed in high-density plots may be associated with the reduced soil pH (Bahram et al. 2018). The primary mechanism leading to lower soil pH at high *A. crenata* local density is unclear. Within high-density plots, specific prokaryotes adapted to low pH environments, particularly the genera *Acidothermus* and *Acidibacter*, were abundant, potentially contributing to organic matter decomposition under acidic conditions (Wang et al. 2019, Oliverio et al. 2020, Ogola et al. 2021, Li et al. 2023). Among the prokaryotic genus enriched in high-density plots, *Conexibacter* and *Phenylobacterium* were particularly noteworthy due to their known association with root exudates of *A. crenata*. *Conexibacter* has been shown to accumulate in the rhizosphere in response to ardisiacrispins, which are triterpenoid saponins exuded from the roots of *A. crenata* (Nakamura and Sugiyama 2025). *Phenylobacterium* is known to be involved in the degradation of triterpenoid saponins (Zhang et al. 2020b), and its relative abundance has been reported to increase with the addition of triterpenoid saponins to the soil (Nakayasu et al. 2021).

While it is challenging to estimate causal relationships in an observational study, our results indicate that the local density of *A. crenata* was, at least in part, a contributing factor to the observed difference in microbial community, mediated by its secondary metabolites. Such plant specialized metabolites exuded from roots of invasive plants can alter soil microbial communities in ways that promote invasion, for example, promoting the colonization of the arbuscular mycorrhizal fungi (Sheng et al. 2022) or exerting allelopathic effects on mutualistic fungi associated with native plants (Anthony et al. 2020). Future evaluation of the impacts of *Conexibacter* and *Phenylobacterium* on *A. crenata* and other native plants will contribute to elucidating the factors driving invasion success through soil modification.

The fungal community exhibited significant compositional shifts, even between uninvaded and low-density plots. Others have reported that fungal taxa may respond more sensitively to changes in soil C/N ratios and moisture than prokaryotic communities (Suzuki et al. 2009, Kaisermann et al. 2015). We found that fungal communities showed greater variability within the same invasion level compared to the prokaryotic community (Supplementary Figures S3, F4), potentially due to stronger dispersal limitations (Bonito et al. 2014, Schmidt et al. 2014, Brigham et al. 2023). In the high-density plots, characteristic indicator taxa included Basidiomycete yeasts, such as *Saitozyma*, which are commonly found in acidic, well-drained soils (Yurkov 2018), forest soils (Buée et al. 2009, Torres-Cruz et al. 2018), and plant roots (Toju et al. 2018). However, the specific ecological roles of these yeasts remain elusive. In addition, Zygomycota taxa such as *Podila* and *Linnemannia* were detected in high-density plots. These fungi include both saprotrophic and endophytic species, suggesting roles in both litter decomposition and root-associated interactions that could affect host performance (Spatafora et al. 2016). While uncertainty remains as to whether fungal community assemblages are primarily shaped by *A. crenata* density or by spatial heterogeneity, our results highlight the interplay between dispersal constraints and invasion pressure, suggesting joint contributions of neutral and density-dependent processes.

## Conclusion

These results strongly suggest that understory shrub species can significantly alter soil chemistry and microbial communities in a mesic forest stand. They also suggest that the local density of invading plants need to be considered explicitly, as the density effects are non-linear. Strictly speaking, this observational study cannot establish a causal relationship between the change in *A. crenata* density and the soil environment. Yet, the direction of observed changes (e.g., lower soil moisture in high-density plots despite their proximity to a creek) strongly suggests that the greater local density of *A. crenata* is the driver, rather than the consequence, of soil environmental conditions. Indeed, increases of labile litter, as well as secondary metabolites (triterpenoid saponins), are likely mechanistic links to cause changes of microbial communities once *A. crenata* becomes high enough. Future studies involving manipulative experiments in a greenhouse that control *A. crenata* density will be promising for elucidating the mechanistic link through which the microbial community responds to the local density of *A. crenata*, possibly leading to positive plant-soil feedbacks that reinforce its invasive dominance.

## Acknowledgments

We would like to extend our gratitude to Betty Schelske for allowing us to use her private property as a sampling site. Chris H Wilson gave critical assistance in plant species identification during the fieldwork. Hiroaki Fujita gave technical support for bioinformatics. Kohmei Kadowaki gave statistical and logical advice and helped improve the manuscript. This research was supported by grants from The Kyoto University Foundation and JSPS Overseas Challenge Program for Young Researchers.

## Author Contributions

Naoto Nakamura was involved in conceptualization (lead), investigation (lead), bioinformatics (lead), validation (equal), writing—original draft (lead), and writing—review and editing (lead). Hirokazu Toju was involved in bioinformatics (equal) and writing—review and editing (equal). S. Luke Flory was involved in conceptualization (equal) and investigation (equal) and writing—review and editing (equal). Kaoru Kitajima was involved in conceptualization (equal) and investigation (equal) and writing—review and editing (equal).

