## Supplementary Figure for "High density invasion by the shrub *Ardisia crenata* in Florida, USA is associated with altered soil chemical properties and microbial communities"

Supplemental Files

| | |
|---|---|
| **Table S1** | Microbial community difference across invasion levels (PerMANOVA) |
| **Figure S1** | Species accumulation (rarefaction) curves of OTUs |
| **Figure S2** | The relationship of soil properties and the local density of *A. crenata* |
| **Figure S3** | The order-level taxonomic composition of microbiome |

**Supplementary Table S1**

Microbial community difference among the three invasion levels (uninvaded, low-density and high-density) tested by PerMANOVA, and statistical difference in heterogeneity of variance in microbial communities tested by Permdisp. Additionally, the results of pair-wise contrast are also shown. Significant differences after Bonferroni corrections (p < 0.017) are shown in bold.

| | Variables | PerMANOVA results | | | PERMDISP results | | |
|---|---|---|---|---|---|---|---|
| | | df | *F* | *P* | df | *F* | *P* |
| Procaryotes | Invasion levels | **2,66** | **30.32** | **< 0.001** | 2,66 | 2.77 | 0.067 |
| | uninvaded vs. low-density | 1,46 | 2.16 | 0.023 | 1,46 | 0.27 | 0.608 |
| | low- vs. high-density | 1,44 | 42.64 | **< 0.001** | 1,44 | 5.03 | 0.028 |
| | uninvaded vs. high-density | 1,42 | 43.37 | **< 0.001** | 1,42 | 2.59 | 0.118 |
| Fungi | Invasion levels | **2,63** | **7.37** | **< 0.001** | 2 ,63 | 0.24 | 0.779 |
| | uninvaded vs. low-density | **1,43** | **1.57** | **0.010** | 1,43 | 0.65 | 0.417 |
| | low- vs. high-density | **1,43** | **10.85** | **< 0.001** | 1,43 | 0.09 | 0.765 |
| | uninvaded vs. high-density | **1,40** | **9.82** | **< 0.001** | 1,40 | 0.12 | 0.734 |

### Supplementary Information

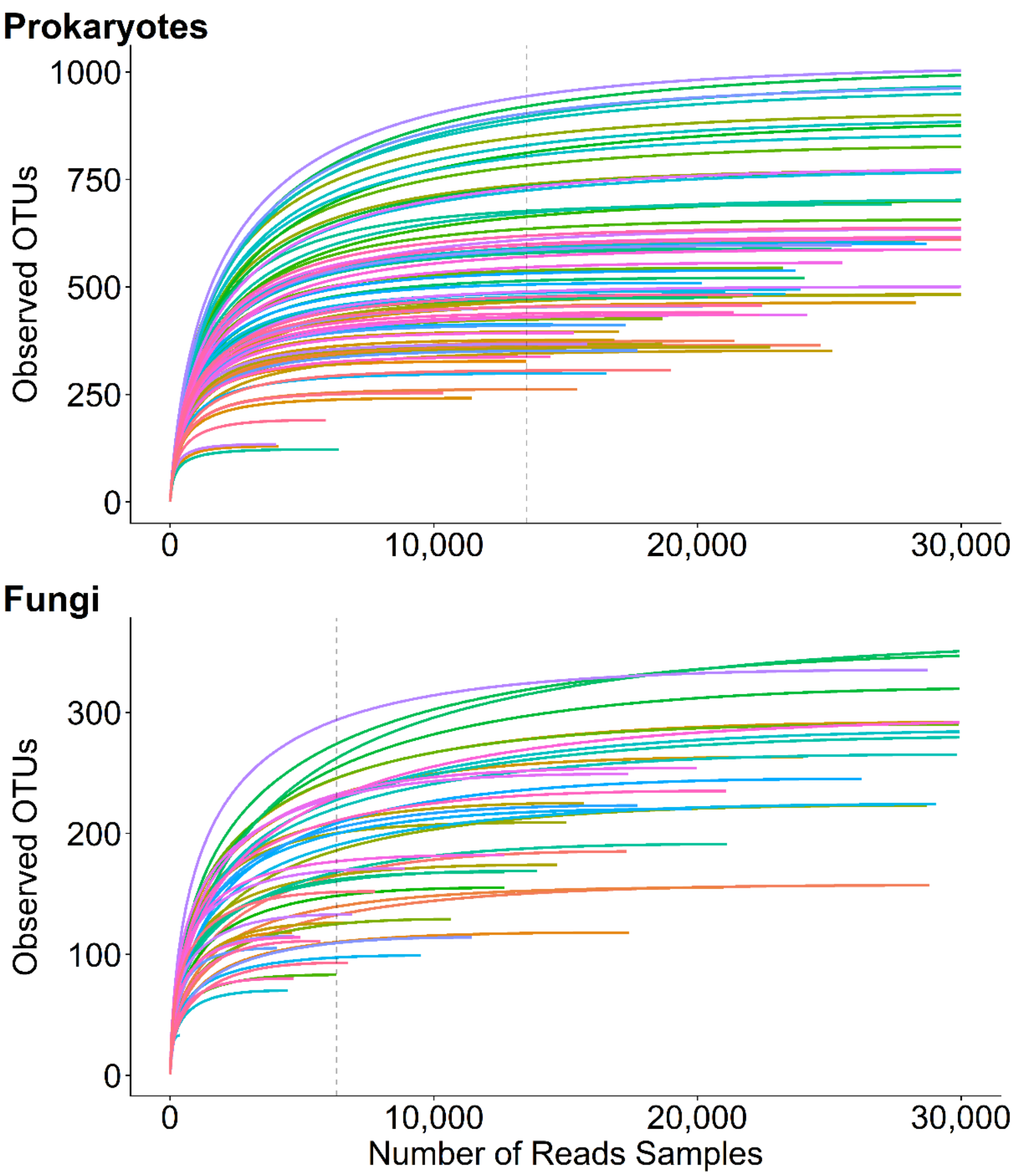


**Supplementary Figure S1** Species accumulation (rarefaction) curves of OTUs detected from 16S rRNA (procaryotes) and ITS (fungi) from a total of 75 soil samples. Each curve represents a soil sample. Vertical dotted lines indicate the sample size to rarefy for the analysis: 13,521 reads for 16S rRNA and 6,314 reads for ITS regions.

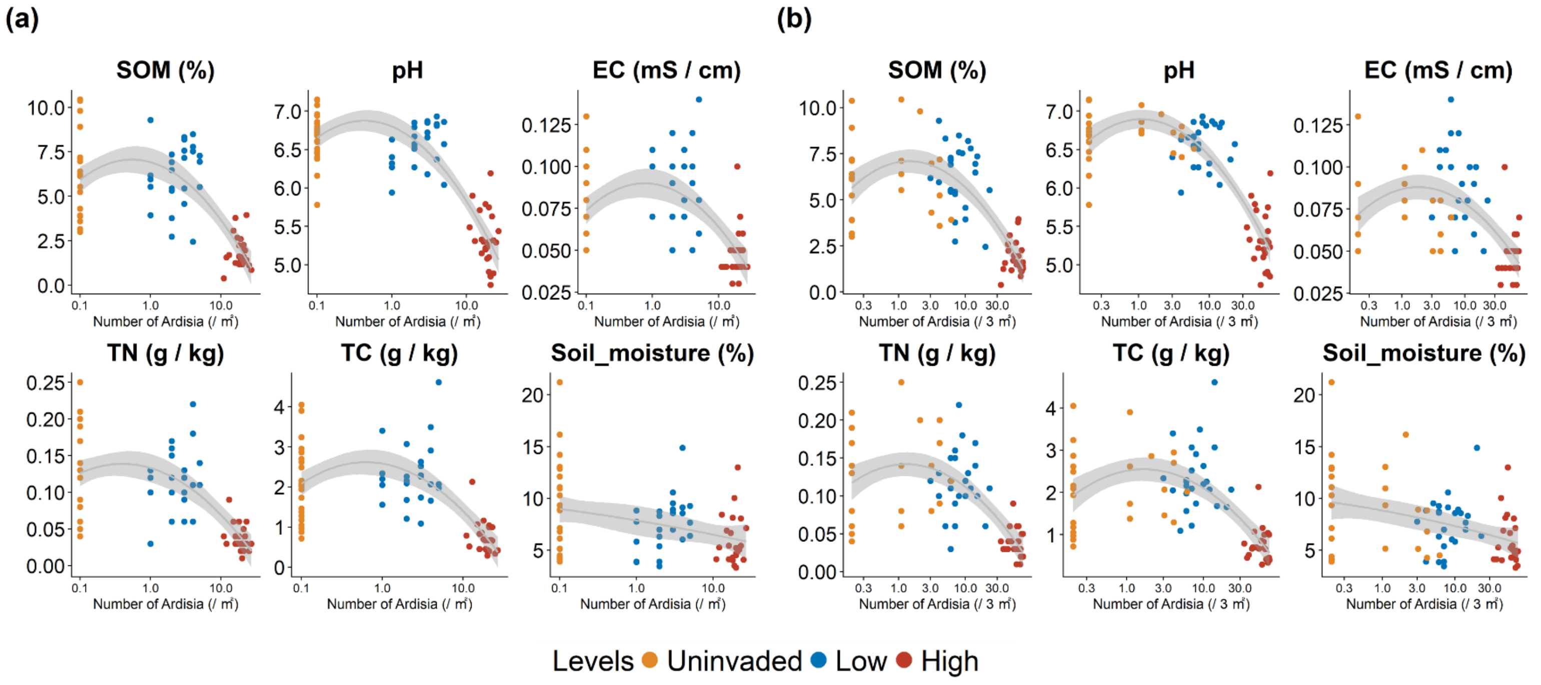


**Supplementary Figure S2**

The relationship of soil properties and the local stem density of *A. crenata* (the number of adult *A. crenata* individuals per square meter, horizontal axis) in the 1-$m^2$ circular area (a) and the 3-$m^2$ circular area (b). Colors represent high-density (reddish brown), low-density (blue), and uninvaded (yellow). The horizontal axis (number of *A. crenata*) was log 10 transformed, and data with zero individuals was changed from zero to 0.1 to accommodate the log transformation. The quadratic regression curve is displayed using the "lm" function of R. (formula = y ~ x + I(x^2)). SOM: soil organic matter; EC: electric conductivity; TN: total nitrogen; TC: total carbon; Soil moisture; soil moisture content.

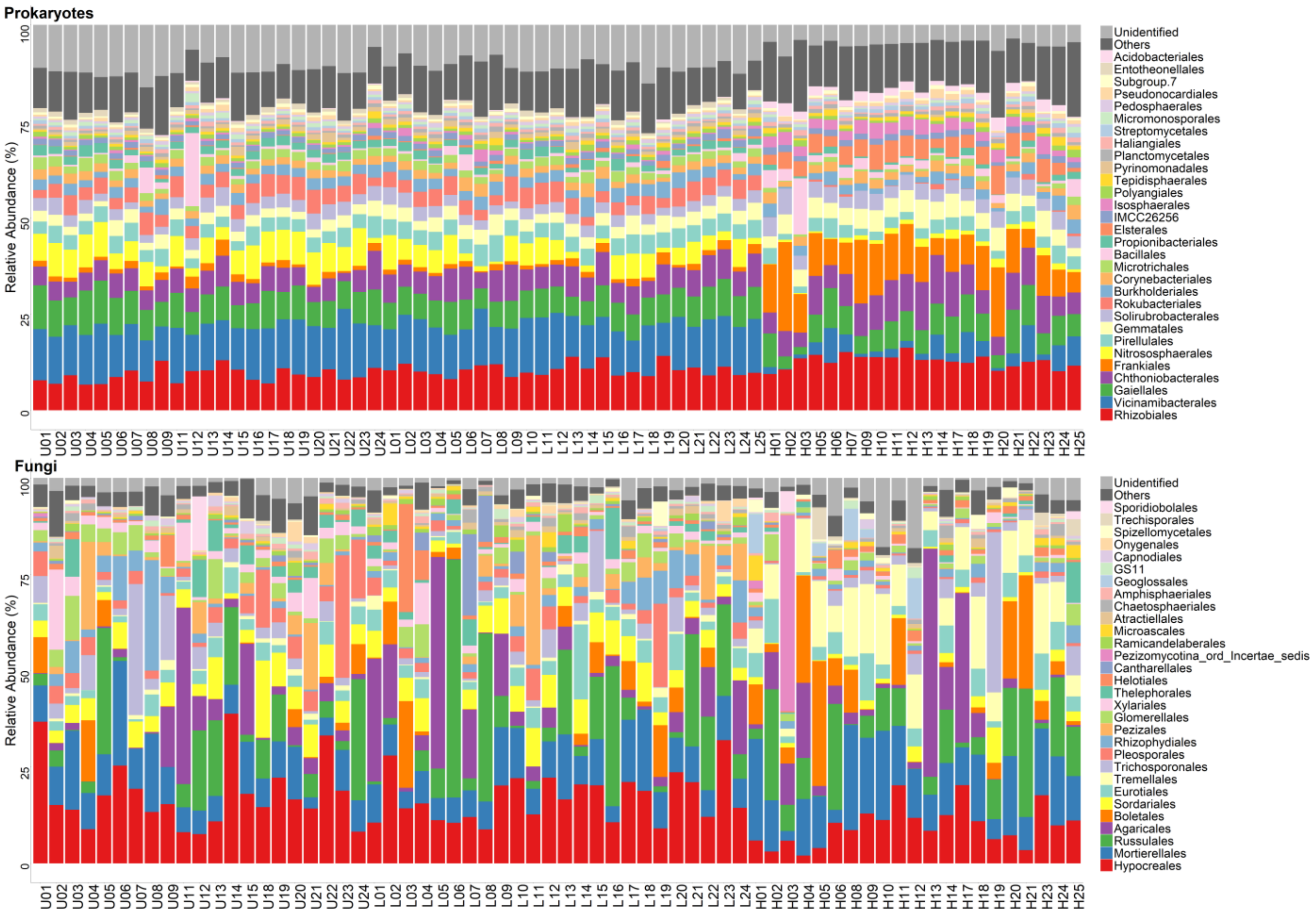


**Supplementary Figure S3** The order-level taxonomic composition of procaryotic and fungal OTUs in each 1-m$^2$ plot. The plot ID starting with H, L, U refers to the level of local density, high-density, low-density, and uninvaded, respectively. The top 30 taxa were displayed. All remaining species are consolidated into the 'Others' category. The vertical axis indicates the relative abundance of each taxon.
